# Estrogen Receptor α Regulates Ethanol Excitation of Ventral Tegmental Area Neurons and Binge Drinking in Female Mice

**DOI:** 10.1101/800052

**Authors:** Bertha J. Vandegrift, Elisa R. Hilderbrand, Rosalba Satta, Rex Tai, Donghong He, Chang You, Cassandre Coles, Hu Chen, Mark S. Brodie, Amy W. Lasek

**Affiliations:** Center for Alcohol Research in Epigenetics and Department of Psychiatry, University of Illinois at Chicago; Department of Physiology and Biophysics, University of Illinois at Chicago, Chicago, Illinois, 60612

## Abstract

Elevations in estrogen (17β-estradiol, E2) are associated with increased alcohol drinking by women and experimentally in rodents. E2 alters the activity of the dopamine system, including the ventral tegmental area (VTA) and its projection targets, which plays an important role in binge drinking. A previous study demonstrated that during high E2 states, VTA dopamine neurons in female mice are more sensitive to ethanol excitation. However, the mechanisms responsible for the ability of E2 to enhance ethanol sensitivity of VTA dopamine neurons have not been investigated. In this study, we used selective agonists and antagonists to examine the role of estrogen receptor subtypes (ERα and ERβ) in regulating the ethanol sensitivity of VTA dopamine neurons and found that ERα promotes the enhanced ethanol response of VTA dopamine neurons. We also demonstrated that the E2-induced increase in ethanol excitation requires the activity of the metabotropic glutamate receptor, mGluR1, which is known to couple with ERα at the plasma membrane. To investigate the behavioral relevance of these findings, we administered lentivirus expressing short hairpin RNAs targeting either ERα or ERβ into the VTA and found that knockdown of each receptor in the VTA reduced binge-like ethanol drinking in female, but not male, mice. Reducing ERα in the VTA had a more dramatic effect on binge-like drinking than reducing ERβ, consistent with the ability of ERα to alter ethanol sensitivity of dopamine neurons. These results provide important insight into sex-specific mechanisms that drive excessive alcohol drinking.

**Significance Statement:** Estrogen has potent effects on the dopamine system and increases the vulnerability of females to develop addiction to substances such as cocaine and alcohol. We investigated the mechanisms by which estrogen increases the response of dopamine neurons in the ventral tegmental area to ethanol. We found that activation of the estrogen receptor, ERα, increased the ethanol-induced excitation of dopamine neurons and that this required the metabotropic glutamate receptor mGluR1. We also demonstrated that estrogen receptors in the ventral tegmental area regulate binge-like alcohol drinking by female, but not male, mice. The influence of estrogen receptors on binge drinking in female mice suggests that treatments for alcohol use disorder in women may need to account for this sex difference.

## Introduction

Binge drinking is defined by the National Institute on Alcohol Abuse and Alcoholism as consuming enough alcohol within a 2-hour period to reach a blood ethanol concentration (BEC) of at least 0.08%. Binge drinking accounts for more than half of the deaths and three-quarters of the economic costs associated with excessive drinking (Stahre et al., 2014; Sacks et al., 2015). Women are more susceptible than men to the devastating health effects associated with alcohol abuse, including liver disease, cardiomyopathy, brain damage, and heightened risk for breast cancer (Agabio et al., 2016; White et al., 2017; Szabo, 2018). Women who binge drink also report having more physically and mentally poor days compared with male binge drinkers (Wen et al., 2012). Unfortunately, more women are drinking excessively now compared with previous decades (Grant et al., 2017). The biological factors that contribute to binge drinking by women are not well understood, although ovarian hormones, specifically estrogen (17β-estradiol, E2), may play a role. Circulating E2 levels in women are positively associated with alcohol consumption (Muti et al., 1998; Martin et al., 1999; Martel et al., 2017), and numerous alcohol drinking studies in female rodents have demonstrated that E2 administration increases alcohol drinking (Ford et al., 2002a; Marinelli et al., 2003; Reid et al., 2003; Ford et al., 2004; Quirarte et al., 2007; Rajasingh et al., 2007; Satta et al., 2018a). Understanding the molecular and cellular mechanisms of action of E2 in enhancing ethanol drinking is important for developing new approaches to reduce excessive drinking by women.

The mesocorticolimbic dopamine (DA) system is critical for the rewarding and reinforcing effects of ethanol (Gonzales et al., 2004; Lovinger and Alvarez, 2017). E2 has potent modulatory effects on this system (Yoest et al., 2018a). For example, E2 enhances cocaine-, amphetamine-, and potassium-stimulated DA release in the striatum of rats and mice (Becker, 1990a, b; Thompson and Moss, 1994; Tobiansky et al., 2016; Yoest et al., 2018b), and potentiates ethanol-stimulated DA release in the prefrontal cortex of female rats (Dazzi et al., 2007). In accord with this finding, ethanol-induced excitation of ventral tegmental area (VTA) DA neurons in female mice is augmented when E2 levels are elevated (Vandegrift et al., 2017). However, the specific estrogen receptor(s) that mediate the enhancement of ethanol-stimulated firing of VTA DA neurons is currently not known.

The first goal of this study was to determine whether estrogen receptor α (ERα) or estrogen receptor β (ERβ) is responsible for the effect of E2 on the response of VTA DA neurons to ethanol. We used selective agonists and antagonists to ERα and ERβ combined with extracellular recordings of VTA DA neurons. Because membrane-bound estrogen receptors can rapidly activate cell signaling pathways through interactions with metabotropic glutamate receptors (mGluRs) and affect neurotransmission and behavior (Tonn Eisinger et al., 2018), the second goal of this study was to determine if the enhancement of ethanol-stimulated VTA DA neuron firing by E2 is dependent on mGluR1 activity.

The third goal of this study was to examine the potential relevance of estrogen receptors expressed in the VTA on alcohol drinking. We used virus-expressed short hairpin (sh)RNAs to reduce the expression of ERα or ERβ in the mouse VTA and measured binge ethanol drinking. Together, the results from our behavioral and electrophysiological experiments indicate that ERα activation in the VTA enhances both ethanol-stimulated firing of VTA DA neurons and binge drinking in female mice. These studies provide important mechanistic and behavioral insights into estrogen receptor signaling in the brain that are relevant to AUD in females.

## Materials and Methods

### Animals

Female C57BL/6J mice were purchased from the Jackson Laboratory (Bar Harbor, ME, USA) at the age of 8 weeks and used for immunohistochemistry, electrophysiology, and behavioral experiments at the age of 10-14 weeks. Transgenic mice containing a bacterial artificial chromosome (BAC) expressing *EGFP* under the control of the *Esr2* promoter were obtained from the Mutant Mouse Regional Resource Center at the University of California, Davis (strain B6.FVB(Cg)-Tg(Esr2-EGFP)ID169Gsat/TmilMmucd, stock 036904-UCD) (Milner et al., 2010) and were used as a reporter for ERβ expression in immunohistochemistry experiments due to a lack of commercially available antibodies specific for mouse ERβ (Snyder et al., 2010). Mice were housed in groups of 3-5 in a temperature- and humidity-controlled room with a 12-hour light/dark cycle (lights on at 6 am), with food and water available *ad libitum*, except water was not available during the limited-access ethanol drinking tests described below. Animal care adhered to the National Institutes of Health *Guide for the Care and Use of Laboratory Animals,* and all procedures were approved by the UIC Animal Care Committee.

### Vaginal cytology

The estrous cycles of gonadally-intact female mice were assessed by vaginal cytology for at least two weeks prior to performing experiments, as previously described (Vandegrift et al., 2017). Briefly, a cotton swab was moistened with sterile water and gently rotated at the vaginal opening. The swab was wiped on a microscope slide, and the smear was immediately analyzed by bright field microscopy using an EVOS® FL inverted microscope (Thermo Fisher Scientific). Estrus was identified by a large quantity of cornified epithelial cells, while diestrus was determined by a predominance of leukocytes (Nelson et al., 1982). These two phases differ in circulating E2 levels. Serum E2 levels in mouse are higher during late diestrus (diestrus II) than estrus and are not statistically significantly different from levels during proestrus as measured by gas chromatography-tandem mass spectrometry (Nilsson et al., 2015).

### Fluorescent immunohistochemistry (IHC)

Mice were euthanized using a lethal dose of a commercial euthanasia solution containing pentobarbital (Somnasol) and transcardially perfused with ice-cold phosphate buffered saline (PBS), followed by 4% paraformaldehyde (PFA). Brains were removed and post-fixed in PFA overnight and cryoprotected in 30% sucrose. Serial coronal sections (40 µm-thick) were collected through the VTA. Sections were blocked in 5% normal donkey serum (Jackson Immunoresearch, #017-000-121, RRID: AB_2337258) and then incubated with various combinations of the following primary antibodies: ERα, rabbit polyclonal, Millipore Sigma, #06-935, RRID: AB_310305; GFP, mouse monoclonal 3E6, Thermo Scientific, #A-11120, RRID: AB_221568; tyrosine hydroxylase (TH), mouse monoclonal LNC1, Millipore Sigma, #MAB318, RRID: AB_827536; TH, rabbit polyclonal, Millipore Sigma, #AB152, RRID: AB_390204. Secondary antibodies were Alexa Fluor 594-conjugated donkey anti-rabbit (Jackson Immunoresearch, #711-585-152, RRID: AB_2340621) and Alexa Fluor 488-conjugated donkey anti-mouse (Jackson Immunoresearch, #715-545-150, RRID: AB_2340846). Sections were mounted onto slides with Vectashield® mounting medium (Vector Laboratories). Images in Fig. 1 were acquired with a Zeiss LSM 710 laser scanning confocal microscope using a 40X objective (all mice were in estrus). Images from Fig. 6 were acquired using an EVOS FL microscope (Thermo Fisher) using a 4X objective.

**Figure 1.**
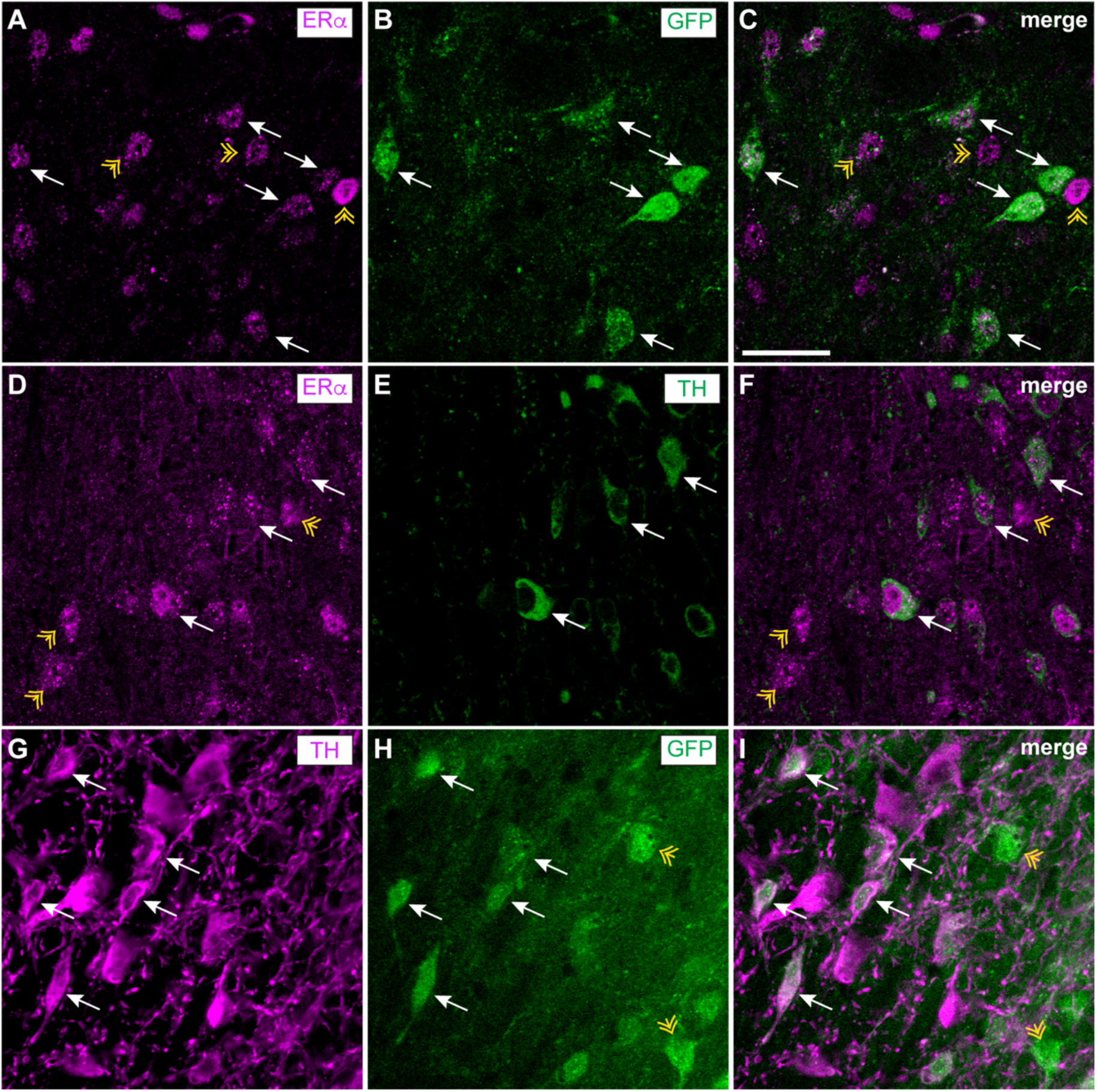
ERα and ERβ are expressed in dopamine neurons and non-dopaminergic cells in the VTA. Brain sections containing the VTA from female mice in estrus were processed for fluorescent IHC with antibodies to ERα, GFP (reporter for ERβ expression), and tyrosine hydroxylase (TH). ***A-C***, Representative images showing ERα (magenta) and GFP (green) colocalization in the VTA of an ERβ-GFP reporter mouse. White arrows indicate examples of ERα+/GFP+ cells, while yellow arrowheads indicate examples of ERα+/GFP-cells. ***D-F***, Representative images showing ERα (magenta) and TH (green) colocalization in the VTA of a C57BL/6J mouse. White arrows indicate examples of ERα+/TH+ cells, while yellow arrowheads indicate examples of ERα+/TH-cells. ***G-H***, Representative images of TH (magenta) and GFP (green) colocalization in the VTA of an ERβ-GFP reporter mouse. White arrows indicate examples of TH+/GFP+ cells, while yellow arrows indicate examples of TH-/GFP+ cells. Scale bar, 50 μm.

### Ovariectomy (OVX)

Mice were anesthetized with intraperitoneal (IP) injections of ketamine (100 mg/kg) and xylazine (8 mg/kg). Hair on the dorsal flanks was trimmed using a hair trimmer, and the skin was disinfected using 70% isopropanol wipes. Bilateral incisions were made in the skin, and a small hole was teased in the muscle wall to allow access to the abdomen. The ovaries and associated fat pads were dissected away from the uterine horns. The muscle wall was sutured, and the skin was closed with wound clips. Mice received a subcutaneous (SC) injection of 2 mg/kg meloxicam immediately after surgery and once on the following day for analgesia. Cessation of the estrous cycle was verified by obtaining vaginal smears for a few days after OVX. Mice recovered for 2 weeks prior to beginning drug treatments.

### *In vivo* drug treatments

Diarylpropionitrile (DPN, selective ERβ agonist, Tocris Bioscience), 4,4’,4’’-(4-Propyl-[1H]-pyrazole-1,3,5-triyl)trisphenol (PPT, selective ERα agonist, Tocris Bioscience), and 17β-estradiol-3-benzoate, herein referred to as E2 (MilliporeSigma) were prepared in solutions of 90% sesame oil/10% ethanol (VEH). DPN and PPT were prepared to final concentrations of 0.5 mg/ml. Mice were injected subcutaneously (SC) with ∼1 mg/kg in a 50 µl volume (final ethanol dose from vehicle solution was < 0.2 g/kg). Mice were treated once per day for three days, with the final dose given 1 h prior to euthanasia. For E2 treatment, 0.2 µg E2 in 50 µl volume (∼10 µg/kg) was injected SC on the first 2 days and 1 µg of E2 in 50 µl volume (∼50 µg/kg) on the third day, 1 h prior to euthanasia. These doses and timing of E2 treatment were chosen because they result in E2 plasma levels similar to proestrus (Vandegrift et al., 2017), when E2 levels peak. The concentrations of PPT and DPN were chosen to balance receptor selectivity with occupancy (Sepehr et al., 2012; Hilderbrand and Lasek, 2018).

### Extracellular recordings and *in vitro* drug treatments

VTA-containing brain slices were prepared for electrophysiology as previously described (Brodie et al., 1999; Dutton et al., 2017). DA VTA neurons were identified both anatomically and based on electrophysiological characteristics that have been well established in the literature and in the Brodie lab (Brodie et al., 1999). Spontaneous spike frequency (firing rate) was recorded and changes in firing rate were determined. A calibrated infusion pump was used to apply ethanol and the mGluR1 antagonist JNJ 16259685 to the aCSF from stock solutions prepared at 100-1000X the final concentration. JNJ 16259685 was prepared in 100% DMSO and the final concentration in the bath was 0.2 µM. The ERβ antagonist PHTPP (4-[2-Phenyl-5,7-*bis*(trifluoromethyl)pyrazolo[1,5-*a*]pyrimidin-3-yl]phenol, Tocris Bioscience) and ERα antagonist MPP (1,3-*Bis*(4-hydroxyphenyl)-4-methyl-5-[4-(2-piperidinylethoxy)phenol]-1*H*-pyrazole dihydrochloride, Tocris Bioscience) were administered via the micropipette by adding the drug to the microelectrode filling solution. Drugs were added to the 0.9% NaCl microelectrode filling solution and time was permitted for the antagonists to diffuse from the recording pipette into the extracellular space around the neuron being recorded.

### Lentiviral vectors

Lentiviruses expressing short hairpin RNAs (shRNAs) targeting *Esr1* (shEsr1-1785, GGCATGGAGCATCTCTACA), *Esr2* (shEsr2-1089, GTACGAAGACAGAGAAGTG), or a sequence not predicted to target any gene in the mouse genome (shScr) were generated from the pLL3.7 vector (Lasek et al., 2007). The *Esr1*-targeting sequence was obtained from Musatov *et al* (Musatov et al., 2006), who demonstrated an 80% reduction in ERα protein and transcript *in vitro* and complete lack of ERα protein in the hypothalamus after adeno-associated virus injection. We previously validated the shEsr1 lentiviral construct used in this study *in vitro* in Satta *et al* (Satta et al., 2018b) and observed a 70% knockdown of *Esr1* transcript. The lentiviral construct expressing shEsr2 was tested for the ability to knockdown *Esr2* in Neuro-2a cells and reduced *Esr2* transcript levels by 94.3 ± 1.5%. We were unable to validate the shEsr2 construct *in vivo* because transcript levels in the VTA are below the limit of detection by qPCR (Cq values ∼34) and because there are no commercially available antibodies specific to mouse ERβ (Snyder et al., 2010). The pLL3.7 vector included a cytomegalovirus (CMV)-enhanced green fluorescent protein (EGFP) reporter cassette for infection site verification after the completion of behavioral testing. Lentiviral plasmids are available from Addgene (#120720 and #120722).

### Stereotaxic injection of lentivirus into the VTA

Gonadally intact, 8- to 12-week-old female C57BL/6J mice were anesthetized with a solution of ketamine and xylazine and placed in a digital stereotaxic alignment apparatus. After bregma alignment and skull leveling, 0.28 mm diameter holes were drilled bilaterally (A/P: −3.2, M/L: ±0.5) for lentivirus microinjection. Lentivirus was delivered to the VTA (D/V: −4.7) at a rate of 0.2 μl/min for a total injection volume of 1 μl/hemisphere. After surgery, mice were maintained on a normal 12-hour light/dark cycle for 1 week. Mice were then transferred to a reversed light/dark cycle room (lights off at 10 am and on at 10 pm) and housed individually for two weeks to acclimate to the change in light/dark cycle before testing drinking.

### Ethanol and sucrose drinking tests

The drinking in the dark (DID) procedure was employed as a model of binge ethanol consumption because it allows mice to achieve blood ethanol levels exceeding 80 mg/dl in a 2-4 h period (Rhodes et al., 2007; Dutton et al., 2017), which is defined as binge drinking by the National Institute on Alcohol Abuse and Alcoholism. For ethanol drinking, mice were given access to a single bottle of 20% ethanol, 3 hours into the dark cycle for 2 h per day on Tuesday-Thursday and 4 h on Friday. Ethanol intake was measured on Friday at 2 and 4 h into the session. Blood was collected from the tail vein immediately after the 4 h session on Friday and blood ethanol concentrations (BECs) were measured using a nicotinamide adenine dinucleotide-alcohol dehydrogenase (NAD-ADH) enzymatic assay (Zapata et al., 2006). Two independent cohorts of mice were injected with lentiviruses and tested for ethanol drinking and data were combined from both cohorts after excluding inaccurate or absent viral infections (shScr, n=10; shEsr1, n=10; shEsr2, n=9). For sucrose drinking, animals were presented with a solution of 2% sucrose in water instead of ethanol using the same DID procedure as described for ethanol. Two independent cohorts of mice were injected with lentiviruses and tested for sucrose drinking and data combined from both cohorts (shScr, n=11; shEsr1, n=10; shEsr2, n=8). Viral infection in the VTA was confirmed for both cohorts, except the in the 2^nd^ shScr (n=5) control group because of an error in tissue processing. However, the two shScr cohorts did not differ statistically in sucrose intake and so the 2^nd^ cohort was included in the data analysis.

### Statistical analysis

Data are presented as the mean ± SEM. Statistical comparisons were made using Student’s *t*-test, one-way ANOVA, two-way ANOVA or two-way repeated measures (RM) ANOVA as indicated for each experiment in the results section. Holm-Sidak’s or Tukey’s multiple comparisons tests were performed as indicated (Prism, Graphpad Software, Inc., La Jolla, CA). A *p* value of < 0.05 was considered significant.

## Results

### ERα and ERβ are expressed in DA neurons and non-DA cells in the mouse VTA

The estrogen receptors, ERα and ERβ, have been detected in the VTA of mice and rats (Kritzer, 1997; Shughrue et al., 1997; Shughrue and Merchenthaler, 2001; Creutz and Kritzer, 2002; Mitra et al., 2003; Kritzer and Creutz, 2008; Milner et al., 2010). In rats, ERα and ERβ immunoreactivity are observed in DA neurons and non-DA cells in the VTA as demonstrated by dual-labeling with an antibody to TH, an enzyme in the DA biosynthetic pathway (Creutz and Kritzer, 2002; Kritzer and Creutz, 2008), but to our knowledge, this has not been investigated in mice. To confirm that ERα and ERβ are present in DA neurons in the mouse VTA, we performed fluorescent IHC on VTA sections of gonadally intact female mice. Transgenic mice expressing EGFP from the *Esr2* promoter (the gene encoding ERβ) were used as a reporter for ERβ expression and the GFP signal was amplified with an antibody to GFP. We first examined GFP and ERα localization and observed both ERα and GFP immunoreactivity in the VTA (Fig. 1*A-C*). Nearly all of the cells expressing GFP were positive for ERα, but there were many cells that were positive for ERα that did not express GFP. These results suggest that ERα and ERβ are co-expressed in some cells in the VTA, but that ERα is expressed more widely than ERβ in the VTA. GFP was visible in TH-positive and TH-negative cells, indicating that ERβ is expressed in both DA and non-DA cells in the mouse VTA (Fig. 1*G-I*). Finally, we examined ERα and TH immunostaining in non-transgenic female C57BL/6J mice and observed that ERα immunoreactivity was visible in both TH-positive and TH-negative cells in the VTA (Fig. 1*D-F*). Together, these results indicate that ERα and ERβ are expressed in DA neurons of the female mouse VTA.

### Baseline firing rates of female mouse VTA DA neurons

A total of 74 VTA neurons from 48 mice were recorded. The initial firing rates of the neurons ranged from 0.54 to 4.66 Hz and the mean firing rate was 1.92 ± 0.11 Hz. There was a significant difference in baseline firing rates between neurons from mice in diestrus and estrus, with a mean firing rate of 2.28 ± 0.25 Hz during estrus and 1.71 ± 0.18 Hz during diestrus (*t*_33_=1.89, *p*=0.034). There were no significant differences in baseline firing rates between neurons from OVX mice treated with VEH (2.34 ± 0.38 Hz), PPT (1.57 ± 0.23 Hz), DPN (1.97 ±0.33 Hz), or E2 (1.85 ± 0.27 Hz; one-way ANOVA, *F*_(2,25)_=1.45, *p*= 0.24).

### Activation of ERα enhances ethanol sensitivity of VTA DA neurons

We previously demonstrated that OVX mice treated with E2 exhibit enhanced ethanol-induced excitation of VTA DA neurons (Vandegrift et al., 2017). In order to determine which estrogen receptor is involved in enhancing ethanol sensitivity, we measured ethanol-induced excitation (40-120 mM) of VTA DA neurons from OVX mice treated with PPT (an ERα-selective agonist), DPN (an ERβ-selective agonist), or VEH. VTA neurons from OVX mice treated with PPT had significantly higher ethanol-induced excitation compared with neurons from DPN- and VEH-treated mice. (Fig. 2, n=10-11 per group, two-way RM ANOVA, ethanol concentration: *F*_(2,56)_ =57.02, *p*<0.0001; treatment: *F*_(2,28)_ = 4.05, *p*=0.028; concentration x treatment interaction: *F*_(4,56)_ =3.06, *p*= 0.024). In response to bath application of 80 mM ethanol, the firing rate of neurons from PPT-treated mice increased by 18.5 ± 1.97%. In comparison, the firing rate of neurons from DPN- and VEH-treated mice increased by 7.7 ± 1.8% and 11.1 ± 2.8%, respectively. Post-hoc Holm-Sidak’s multiple comparisons test demonstrated a significant increase in ethanol-induced excitation in response to 80 mM ethanol from mice treated with PPT compared with VEH (*p*=0.046) and DPN (*p*=0.0044), and in response to 120 mM ethanol from mice treated with PPT compared with VEH (*p*=0.038) and DPN (*p*=0.0082). These results indicate that activation of ERα enhances the response of VTA DA neurons to ethanol in OVX mice and suggests that the E2-induced increase in ethanol sensitivity is likely due to activation of ERα.

**Figure 2.**
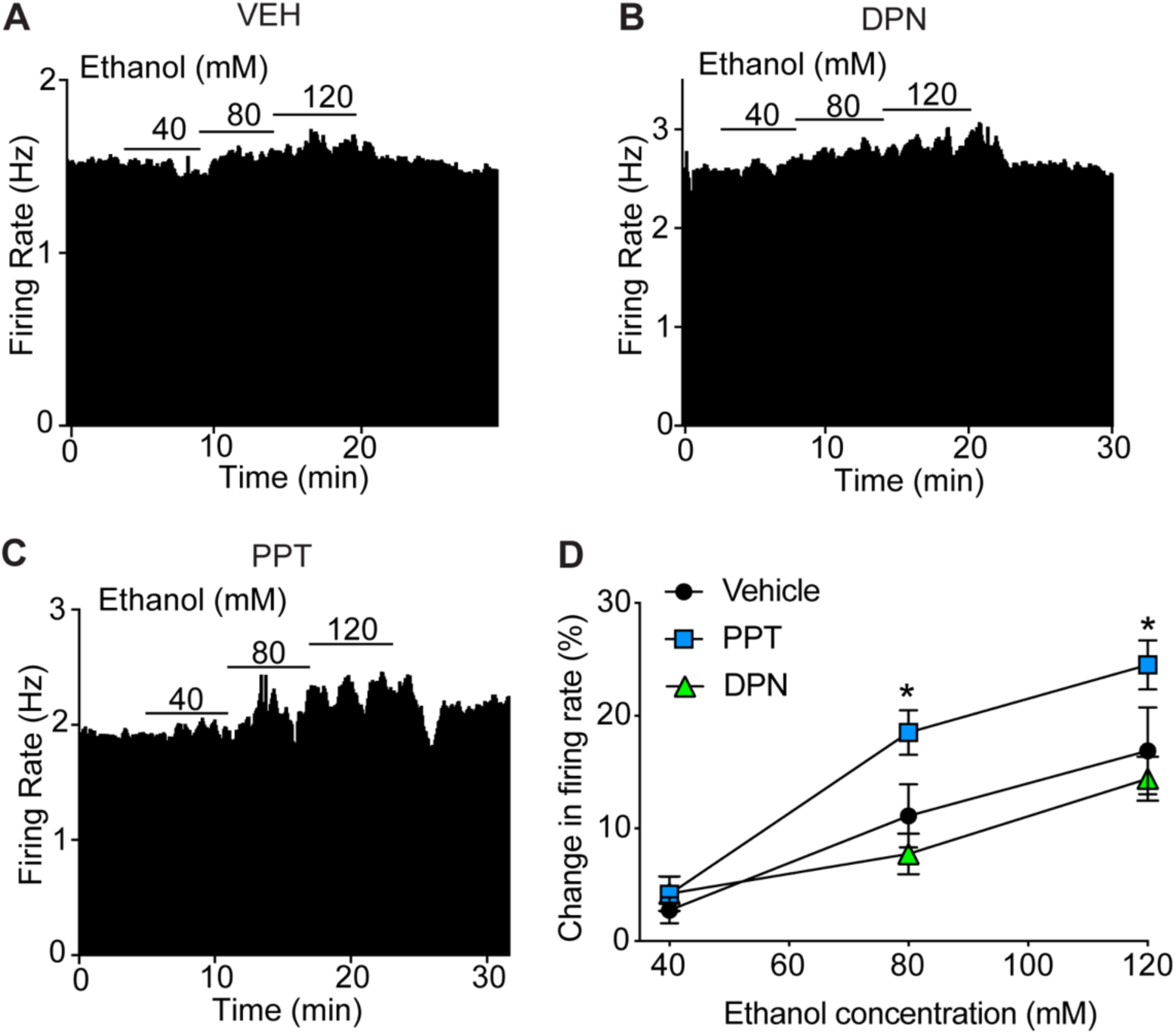
Activation of ERα enhances ethanol sensitivity of VTA DA neurons. Extracellular recordings were obtained from VTA DA neurons in ovariectomized (OVX) mice treated systemically with an ERβ agonist (DPN), ERα agonist (PPT), or vehicle (VEH). ***A-C***, Rate meter graphs showing response to 40-120 mM ethanol of a representative neuron from an OVX mouse treated with VEH (***A***), DPN (***B***), or PPT (***C***). ***D***, Ethanol concentration-response graph (40, 80, and 120 mM). Neurons from OVX mice treated with PPT (n=10), but not DPN (n=10), had enhanced excitation by ethanol compared with VEH-treated mice (n=10). \**p* < 0.05 by post hoc Holm-Sidak’s multiple comparisons test.

### ERα regulates ethanol sensitivity of VTA DA neurons from mice in diestrus

VTA DA neurons from mice in diestrus are more sensitive to ethanol excitation when compared with neurons from mice in estrus (Vandegrift et al., 2017). This enhancement of ethanol-induced excitation in diestrus is blocked by acute administration of the ERα/ERβ antagonist, ICI 182,780, directly to the VTA slice (Vandegrift et al., 2017). To determine if this effect results from blocking the activity of a specific estrogen receptor, we administered MPP, an ERα-selective antagonist, or PHTPP, an ERβ-selective antagonist, to VTA slices from mice in diestrus or estrus using the recording pipette. The response to 80 mM ethanol was tested prior to administration of the antagonist and again 70-80 minutes later, after the antagonist diffused from the recording pipette onto the cell of interest. The ERα antagonist MPP decreased the excitatory response of VTA DA neurons to ethanol from mice in diestrus, but had no effect on neurons from mice in estrus (Fig. 3*A-C*, n = 5-6 per group, two-way RM ANOVA, time: *F*_(1,9)_= 7.46, *p* = 0.023; phase: *F*_(1,9)_ = 3.50, *p* = 0.094; time x phase interaction: *F*_(1,9)_ = 10.50, *p* = 0.01). Post-hoc Holm-Sidak’s multiple comparisons test demonstrated a significant difference in ethanol-induced excitation between estrus (9.9 ± 1.4% increase) and diestrus (15.9 ± 0.8% increase) prior to MPP treatment (*p*=0.0054), consistent with our previous findings (Vandegrift et al., 2017). After administration of MPP, there was no longer a difference (*p*=0.60) in ethanol-induced excitation between diestrus (9.6 ± 1.3% increase) and estrus (10.5 ± 1.1% increase) because of a significant reduction in ethanol-induced excitation in neurons from mice in diestrus (*p*=0.0058). These results suggest that ERα acutely regulates ethanol sensitivity in the VTA in an estrous cycle phase-dependent manner.

**Figure 3.**
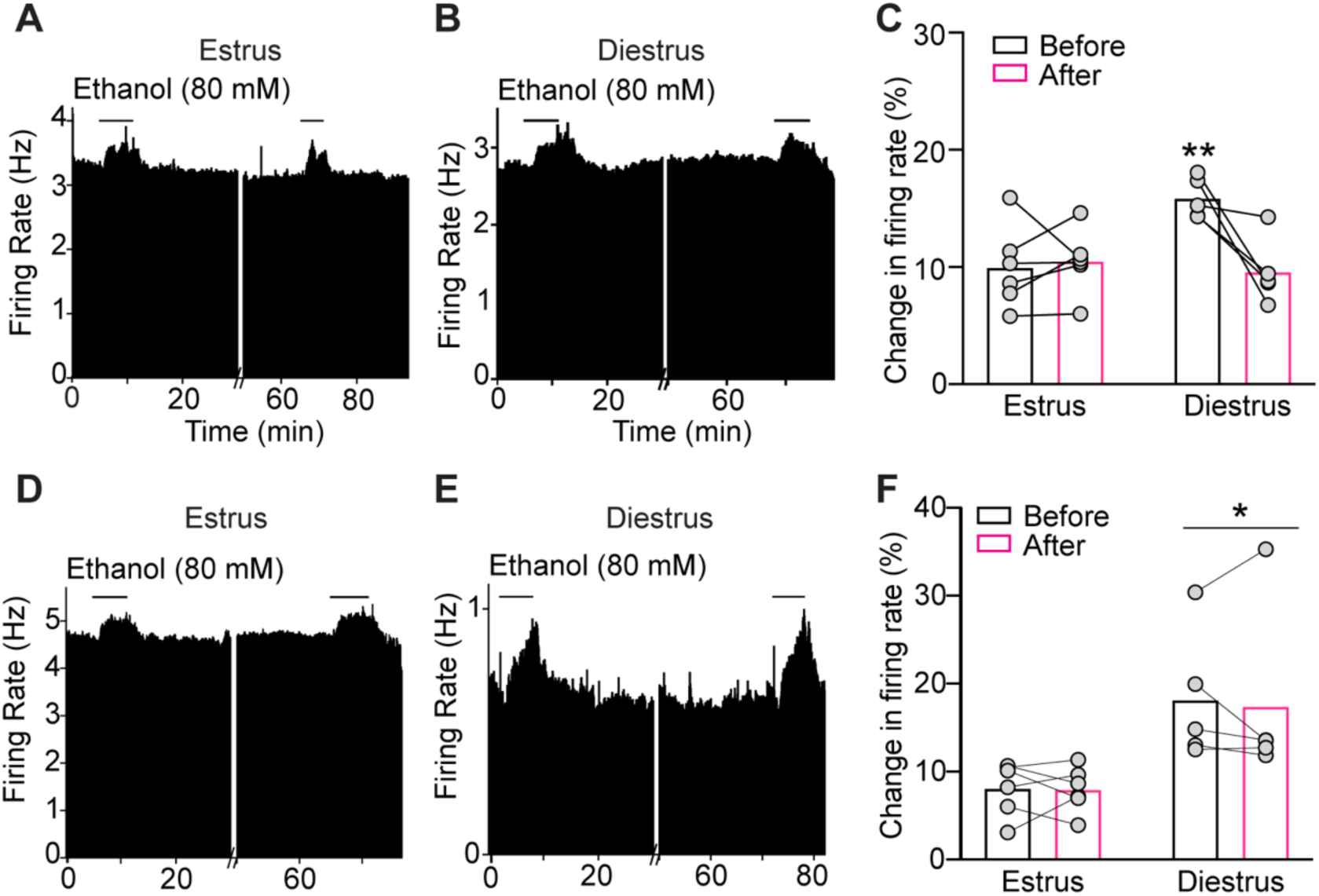
ERα regulates ethanol sensitivity of VTA DA neurons from mice in diestrus. ***A-B,*** Rate meter graphs showing the response of a representative neuron to 80 mM ethanol before and after treatment with the ERα antagonist MPP from mice in estrus (***A***) and diestrus (***B***). ***C***, Graph of neuronal responses to 80 mM ethanol from mice in estrus (n=5) and diestrus (n=6) before and after treatment of slices with MPP. The enhanced ethanol-induced excitation of neurons from mice in diestrus was decreased after treatment with MPP, whereas the ethanol responses of DA VTA neurons from mice in estrus are unchanged by MPP. \*\**p* < 0.05 when comparing diestrus with estrus before MPP treatment and when comparing responses in diestrus before and after MPP by post-hoc Sidak’s multiple comparisons test. ***D-E***, Rate meter graphs showing the response of a representative neuron to 80 mM ethanol before and after treatment with the ERβ antagonist PHTPP from mice in estrus (***D***) and diestrus (***E***). ***F***, Graph of neuronal responses to 80 mM ethanol from mice in estrus (n=6) and diestrus (n=5) before and after treatment of slices with PHTPP. There was no change in ethanol excitation after PHTPP treatment, but there was a significant main effect of cycle phase. \**p*< 0.05, main effect of cycle by two-way ANOVA.

In contrast to results obtained with the ERα antagonist, the ERβ antagonist PHTPP had no effect on ethanol sensitivity of VTA DA neurons. As expected, there was a significant main effect of estrous cycle phase, indicating enhanced ethanol excitation in neurons from mice in diestrus compared with estrus (Fig. 3*D-F*, n = 5-6 per group, two-way RM ANOVA, time: *F*_(1,9)_= 0.18, *p*=0.68; phase: *F*_(1,9)_=7.17, *p*=0.025; time x phase interaction: *F*_(1,9)_ = 0.088, *p*=0.77). Neurons from mice in estrus exhibited no increase in the response to ethanol, as ethanol increased firing by 8.0 ± 1.2% and 7.9 ± 1% before and after PHTPP delivery, respectively. Likewise, while neurons from mice in diestrus initially responded to 80 mM ethanol with an 18.1 ± 3.3% increase in firing rate, after PHTPP delivery, these neurons responded similarly to ethanol with a 17.4 ± 4.5% increase in firing rate. Together, these results indicate that ERα, but not ERβ, enhances the ethanol-induced excitation of VTA DA neurons in female mice during diestrus, when E2 levels are higher than in estrus.

### mGluR1 is required for the E2-induced enhancement of ethanol excitation

E2 rapidly activates membrane-bound estrogen receptors that couple to mGluRs in hippocampal (Huang and Woolley, 2012) and striatal (Grove-Strawser et al., 2010) neurons. We hypothesized that mGluR activation might be occurring in the VTA of mice after E2 treatment and thus contributing to the enhancement of ethanol-simulated firing. To test this, OVX mice were administered E2 or VEH and the response to 40-120 mM ethanol was measured in VTA DA neurons before and after bath application of the mGluR1 antagonist, JNJ 16259685. JNJ 16259685 did not affect ethanol excitation of VTA DA neurons from OVX mice treated with VEH (Fig.4*A-B*, n = 5 per group, two-way RM ANOVA, treatment: *F* _(1, 4)_ = 0.52, *p*=0.51; ethanol concentration: *F*_(2,8)_ = 21.47, *p*=0.0006; treatment x concentration interaction: *F*_(2,8)_ = 1.39, *p* = 0.30). In contrast, JNJ 16259685 decreased the response of VTA DA neurons to ethanol in OVX mice treated with E2 (Fig. *4C-D*, n = 6 per group, two-way RM ANOVA, treatment: *F*_(1,5)_ = 111.58, *p* = 0.019; ethanol concentration: *F*_(2,10)_ = 30.44, *p*<0.0001; treatment x concentration interaction: *F*_(2,10)_ =1.56, *p*=0.26). Neurons from E2-treated OVX mice initially responded to 80 mM ethanol with a 17.8 ± 2.7% increase in firing rate. After 45 minutes of JNJ 16259685 exposure, the neurons responded to ethanol with a 9.9 ± 3.1% increase in firing rate. These results indicate that E2-mediated enhancement of excitation by ethanol depends on mGluR1 activation.

**Figure 4.**
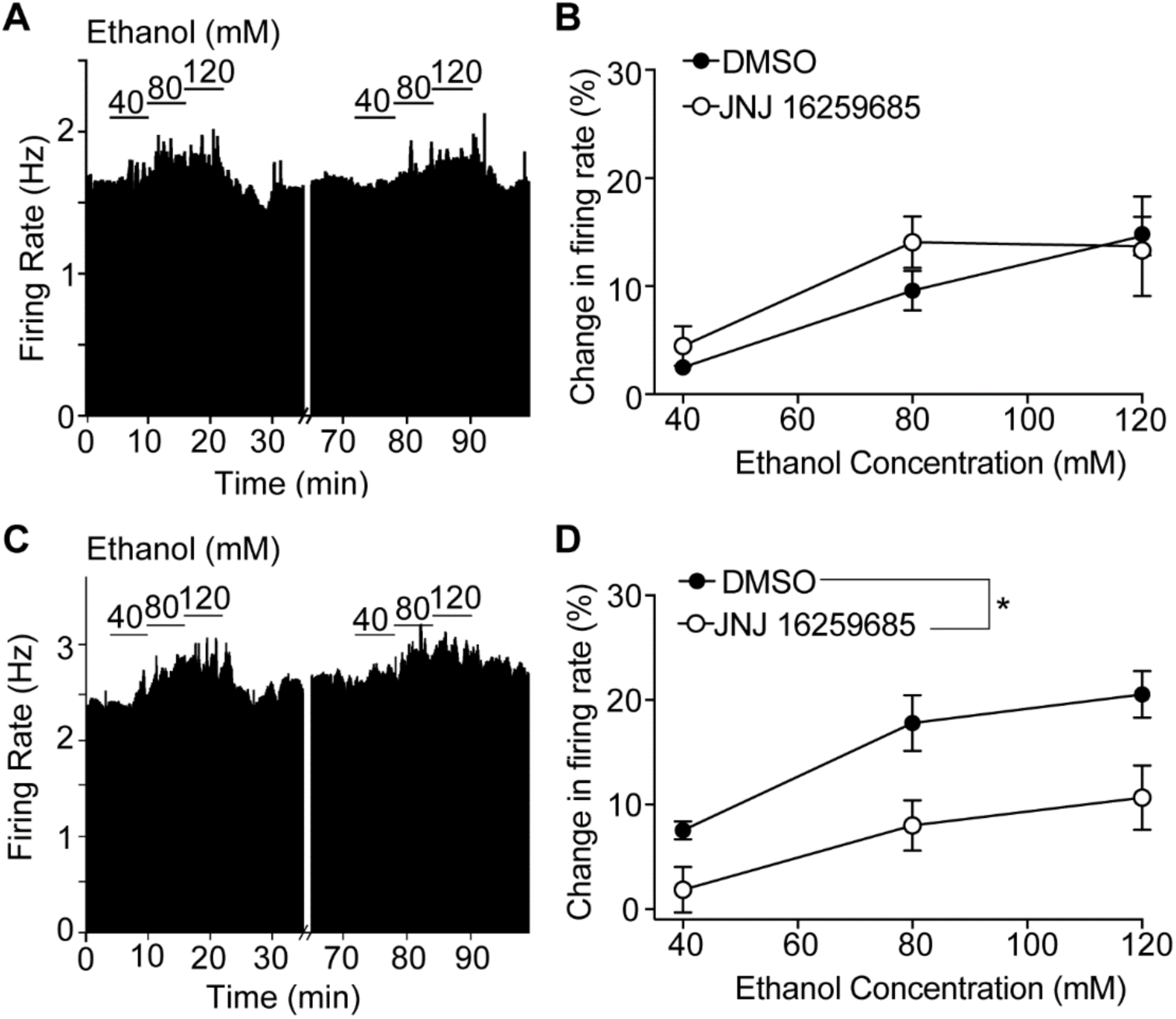
mGluR1 is required for the E2-induced enhancement of ethanol excitation. ***A***, Rate meter graph showing the response to 40-120 mM ethanol of a representative neuron from an ovariectomized (OVX) mouse treated with vehicle before and after treatment with the mGluR1 antagonist JNJ 16259685. ***B***, Ethanol concentration-response graph of neurons (n=5) from OVX mice treated with vehicle showing responses before (DMSO vehicle) and after slice treatment with JNJ 16259685. ***C***, Rate meter graph showing the response to 40-120 mM ethanol of a representative neuron from an E2-treated OVX mouse before and after treatment with JNJ 16259685. ***D***, Ethanol concentration-response graph of neurons (n=6) from E2-treated OVX mice before and after treatment of slices with JNJ 16259685. The ethanol excitation of neurons from E2-treated OVX mice was significantly reduced after JNJ 16259685. \**p* < 0.05, main effect of treatment by two-way RM ANOVA.

### mGluR1 regulates ethanol sensitivity of VTA DA neurons from mice during diestrus

We tested whether the increased ethanol sensitivity we observed during diestrus compared with estrus could be attributed to activation of mGluR. We measured the response to 40-120 mM ethanol in VTA DA neurons from mice in diestrus and estrus before and after bath application of JNJ 16259685. Ethanol excitation of VTA DA neurons from females in estrus did not change after treatment with JNJ 16259685 (Figure 5*A-B*, n = 5 per group, two-way RM ANOVA, treatment: *F*_(1,4)_ = 0.18, *p*=0.70; ethanol concentration: *F*_(2,8)_ = 60.66, *p* < 0.0001; treatment x concentration interaction: *F*_(2,8)_ =1.51, *p*= 0.28). For example, neurons from mice in estrus exhibited a 10.4 ± 1.6% increase in firing in response to 80 mM ethanol before treatment with the mGluR1 antagonist and an 8.9 ± 1.7% increase after treatment. In contrast, treatment with JNJ 16259685 decreased ethanol excitation of VTA DA neurons from females in diestrus (Fig. 5*C-D*, n = 6 per group, two-way RM ANOVA, treatment: *F*_(1,5)_ =7.44, *p*=0.041; ethanol concentration: *F*_(2,10)_ = 6.56, *p*=0.015; treatment x concentration interaction: *F*_(2,10)_ =2.46, *p*=0.14). Neurons from mice in diestrus responded to 80 mM ethanol with a 14.9 ± 1.6% increase in firing rate before treatment with the mGluR1 antagonist and a 7.9 ± 3.1%. increase in firing rate after treatment. These results indicate that mGluR1 activity is required for the increased sensitivity of VTA DA neurons to ethanol from mice in diestrus, similar to what we observed in mice treated with E2.

**Figure 5.**
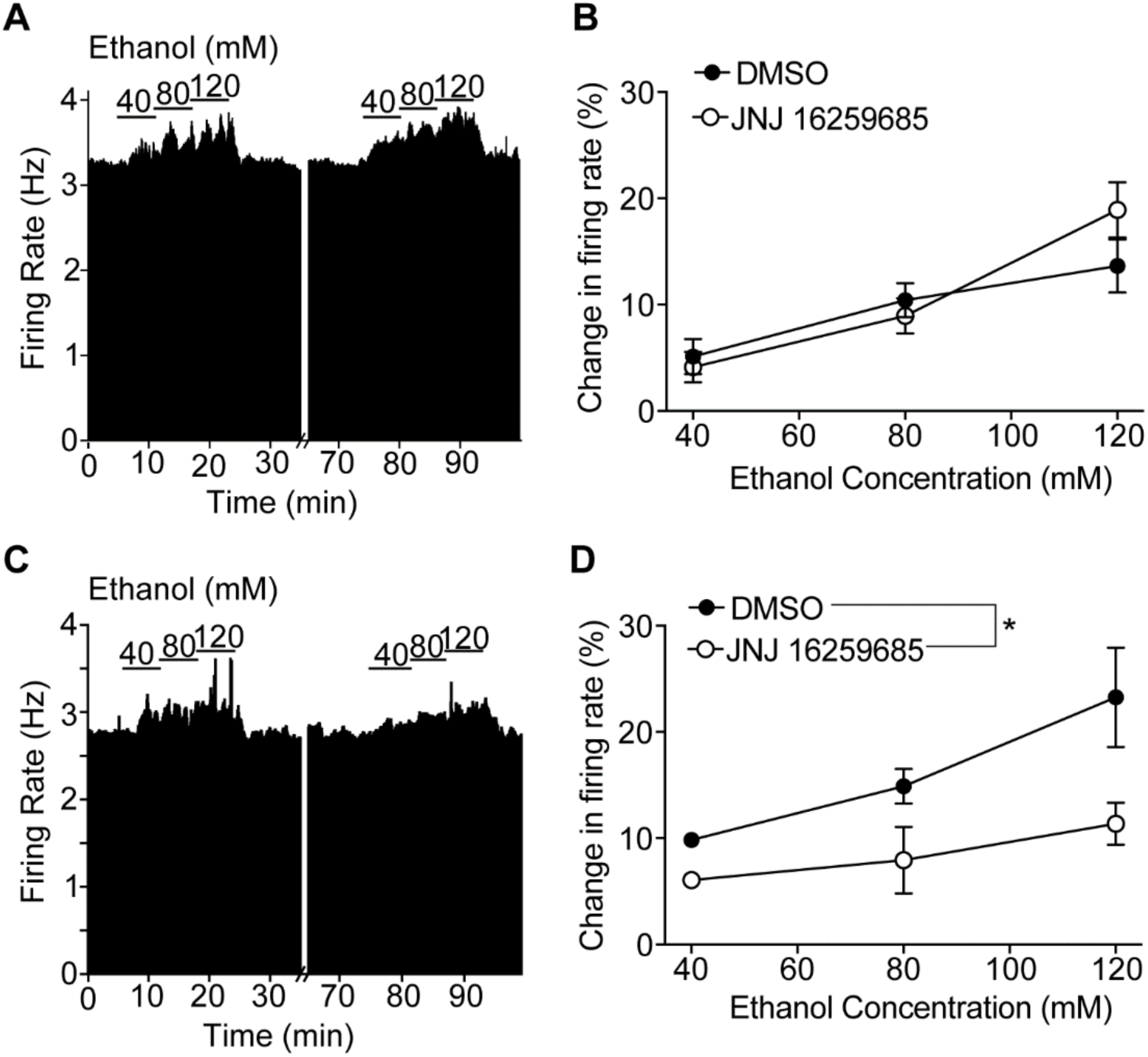
mGluR1 regulates ethanol sensitivity of VTA DA neurons from mice during diestrus. ***A***, Rate meter graph showing the response of an individual neuron to 40-120 mM ethanol before and after treatment with the mGluR1 antagonist JNJ 16259685 from a mouse in estrus. ***B,*** Ethanol concentration-response graph of neurons (n=5) from mice in estrus before and after treatment of slices with JNJ 16259685. ***C***, Rate meter graph showing the response of an individual neuron to 40-120 mM ethanol before and after treatment with JNJ 16259685 from a mouse in diestrus. ***D***, Ethanol concentration-response graph of neurons (n=6) from mice in diestrus before and after treatment of slices with JNJ 16259685. The ethanol excitation of neurons from mice in diestrus was significantly reduced after treatment. *p < 0.05, main effect of treatment by two-way RM ANOVA.

### Intra-VTA knockdown of estrogen receptors reduces binge ethanol consumption in female mice

Female mice drink more alcohol than male mice in a binge drinking test, an effect that is attributed to E2 (Satta et al., 2018a). Our results demonstrating that ERα activation increases the sensitivity of VTA DA neurons to ethanol suggest that ERα expression in the VTA might regulate binge drinking because of the critical role of the VTA in this behavior. To determine if estrogen receptors in the VTA are important for alcohol drinking, we used shRNAs to locally knockdown expression of *Esr1* (encoding ERα) or *Esr2* (encoding ERβ) in the VTA of gonadally intact female mice. Mice were injected with lentiviral vectors expressing shEsr1, shEsr2, or a shScr control and tested for binge ethanol drinking using the drinking in the dark test 3 weeks after injection. Two-way RM ANOVA revealed significant main effects of shRNA and time on ethanol intake during the 2-hour drinking sessions, but no treatment x time interaction (Fig. 6*A*, shRNA: *F*_(2,26)_ = 13.10, *p* < 0.0001; time: *F*_(3,78)_ = 6.02, *p* = 0.001; interaction: *F*_(6,78)_ = 0.61, *p* = 0.72). Post-hoc testing to identify differences in shRNA groups demonstrated that reducing *Esr1* expression in the VTA resulted in a 30% decrease (*p* < 0.0001) in 2-hour ethanol intake over the 4 days. Reducing *Esr2* expression in the VTA also resulted a significant, albeit less pronounced, 16% decrease (*p* = 0.038) in 2-hour ethanol intake over the 4 days. Ethanol intake during the 4-hour session on day 4 was attenuated by a reduction in estrogen receptor levels in the VTA (Fig. 6*B, F* _(2, 26)_ = 6.85, *p* = 0.0041). Multiple comparisons testing revealed a significant 29% decrease in ethanol drinking in mice expressing shEsr1 (*p* = 0.0030) and a non-significant 17% decrease in mice expressing shEsr2 (*p* = 0.097) in the VTA during the 4-hour session. Consistent with reduced ethanol intake during the 4-hour session, BECs were lower in mice with estrogen receptor knockdown in the VTA (Fig. 6*C*, one-way ANOVA, *F*_(2, 25)_ = 3.35, *p* = 0.051). This effect was driven by a 55% reduction in BECs in the shEsr1 treatment group (*p* = 0.041), as BECs in the shEsr2 group were not significantly different from the shScr group (24% reduction, *p* = 0.51). We also tested water intake one day prior to measuring ethanol consumption in the same mice and did not observe an effect of estrogen receptor knockdown on water intake during a 2-hour period (Fig. 6*D*), indicating that reducing estrogen receptors in the VTA does not affect general fluid intake. Viral infection in the VTA was confirmed by dual immunofluorescent staining of brain sections using antibodies to TH and GFP (Fig. 6*G-J*). Selective knockdown of estrogen receptors in the VTA did not affect estrous cycles (data not shown), as mice continued to cycle normally throughout the duration of the experiment. Binge ethanol drinking also did not significantly vary during the estrous cycle, as reported previously (Satta et al., 2018a), nor was drinking altered only at specific phases of the estrous cycle by knockdown of estrogen receptors (data not shown). These results indicate that reducing levels of estrogen receptors in the VTA of female mice decreases binge-like drinking, with a stronger effect of ERα knockdown.

**Figure 6.**
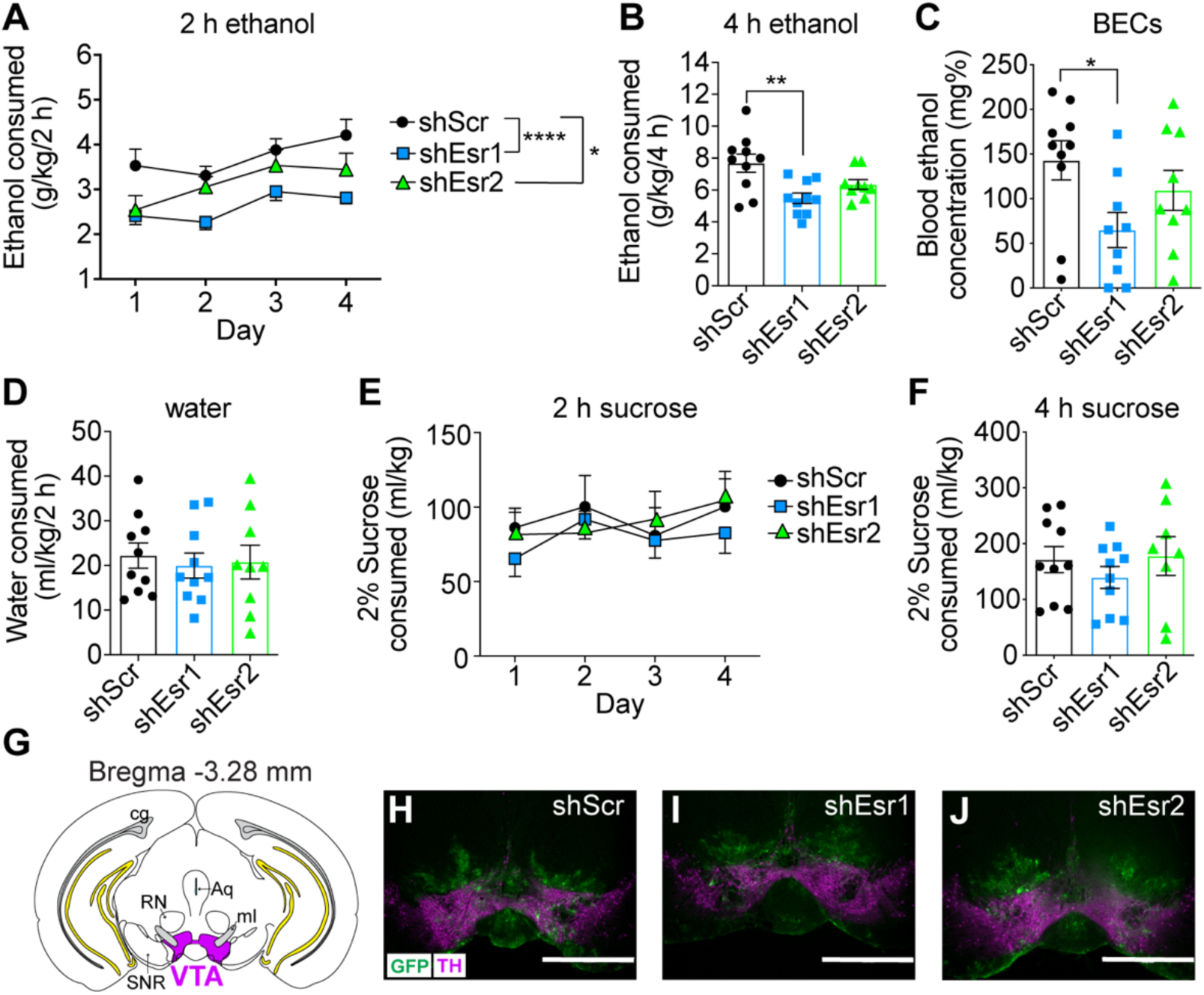
Intra-VTA knockdown of estrogen receptors reduces binge ethanol consumption in female mice. Gonadally intact female mice were injected with lentivirus expressing either shScr (control, n=10), shEsr1 (n=10), or shEsr2 (n=9) and tested for binge ethanol drinking in the drinking in the dark test. ***A,*** Ethanol intake in g/kg over 2 h on each day of drinking. \**p*<0.05, \*\*\*\**p*<0.0001. ***B,*** Ethanol intake in g/kg during the final 4-h drinking session. \*\**p*<0.01. ***C,*** Blood ethanol concentrations (BECs) after the final 4-h drinking session. ***D,*** Water intake during a 2-h session conducted 1 day prior to testing ethanol drinking. ***E,*** Separate groups of mice were injected with lentivirus expressing shScr (n=11), shEsr1 (n=10), or shEsr2 (n=8) and tested for consumption of 2% sucrose using the same procedure as was used for ethanol drinking. Shown is sucrose intake in ml/kg over 2 h on each day of drinking. ***F,*** Sucrose intake in ml/kg during the final 4 h session. ***G,*** Illustration of a coronal brain section showing injection site in the VTA (shaded magenta). For reference, hippocampal structures are colored in yellow and the cingulum is shaded grey. ***H-J,*** Representative images of infection in the VTA of mice injected with lentivirus expressing shScr (***H***), shEsr1 (***I***), or shEsr2 (***J***). GFP is in green and TH is in magenta. Scalebar, 1 mm. Abbreviations, RN, red nucleus; Aq, aqueduct; SNR, substantia nigra pars reticulata; ml, medial lemniscus.

In order to determine if the effects of intra-VTA estrogen receptor knockdown on binge-like drinking were specific to ethanol or might extend to other rewarding substances, we performed the 4-day drinking in the dark test using 2% sucrose (instead of ethanol) with mice expressing shEsr1, shEsr2, or shScr in the VTA. There were no significant effects of shRNA, time, or an shRNA x time interaction for sucrose drinking during the 2-hour sessions or during the final 4-hour session (Fig. 6*E-F*), demonstrating that knockdown of estrogen receptors in the VTA of female mice does not affect consumption of all rewarding substances. Finally, to determine if the effect of estrogen receptor knockdown in the VTA is sex-specific, we injected lentiviruses expressing shScr, shEsr1, or shEsr2 into the VTA of male mice and tested them in the ethanol drinking in the dark test. Despite the fact that estrogen receptors are expressed in the VTA of males, knockdown of *Esr1* or *Esr2* did not alter ethanol drinking in male mice (Fig. 6-1). Fluorescent immunostaining with antibodies to GFP and ERα demonstrated a 54% reduction in ERα levels in shEsr1virus-infected cells. These results suggest a sexually dimorphic role for estrogen receptors in the VTA in ethanol binge drinking.

## Discussion

The primary novel findings of this study are that: 1) activation of ERα potentiates ethanol-induced excitation of VTA DA neurons, and 2) ERα, and to a lesser extent ERβ, in the VTA regulates binge-like ethanol drinking. We previously demonstrated that VTA DA neurons from mice in diestrus and E2-treated OVX mice are more sensitive to ethanol excitation than neurons from mice in estrus and VEH-treated OVX mice (Vandegrift et al., 2017). Multiple studies have found that E2-treated females consume more ethanol under various access conditions, including those that promote binge drinking (Ford et al., 2002a; Marinelli et al., 2003; Reid et al., 2003; Ford et al., 2004; Quirarte et al., 2007; Rajasingh et al., 2007; Satta et al., 2018a). Prior to this study, it was not known which estrogen receptor(s) are responsible for these behavioral and neurophysiological effects.

Evidence that ERα is responsible for the increased sensitivity of VTA DA neurons to ethanol in females is provided by our experiments in OVX mice treated with the selective ERα agonist, PPT, and in brain slices from mice in diestrus treated with the ERα antagonist, MPP. PPT treatment of OVX mice mimicked the effect of E2 treatment, indicating that activation of ERα is most likely the mechanism by which E2 increased ethanol excitation of VTA DA neurons. We confirmed a role for ERα in the enhanced sensitivity to ethanol in gonadally-intact mice in diestrus, showing that acute application of MPP decreased ethanol induced excitation when E2 levels are rising (Nilsson et al., 2015). This result demonstrates that ERα is important for the enhanced response to ethanol in a natural hormonal state, and not just in gonadectomized mice treated with E2. In contrast to the results obtained with ERα, pharmacological manipulation of ERβ did not alter the potency of ethanol on VTA DA neurons, even though ERβ is expressed in these cells. Together, these data demonstrate an important role for E2 acting on ERα to increase ethanol sensitivity of VTA DA neurons. Although we showed that ERα is expressed in both DA and non-DA cells in the VTA, the ERα-mediated enhancement of ethanol sensitivity could conceivably be due to ERα expression on neuronal terminals in the VTA, such as those from the medial preoptic area that project to the VTA and that are known to regulate DA release in the nucleus accumbens (Tobiansky et al., 2016; McHenry et al., 2017). Further investigation is needed to identify the specific cell types in which ERα regulates responses to ethanol in the VTA.

Our results suggest that activation of ERα would increase ethanol-stimulated DA release in target regions such as the nucleus accumbens, which might contribute to enhanced rewarding and reinforcing effects of ethanol during high-estrogen states. Dazzi *et al* demonstrated that ethanol increased DA release in the prefrontal cortex of OVX rats treated with E2, but not in untreated OVX rats (Dazzi et al., 2007), and that treatment with the selective estrogen receptor modulator clomifene prevented the ethanol-induced DA elevation. It remains to be determined whether treatment with an ERα-selective agonist or antagonist would alter ethanol-induced elevation of DA in regions such as the nucleus accumbens and prefrontal cortex.

It is important to point out that basal firing rates of VTA DA neurons from mice in estrus were slightly higher than neurons from mice in diestrus. Other groups have shown that basal firing rate and bursting of VTA neurons are greater in estrus compared with diestrus (Zhang et al., 2008; Calipari et al., 2017). Supporting this observation, DA levels in the prefrontal cortex and striatum of rats are also higher in estrus compared with diestrus (Xiao and Becker, 1994; Dazzi et al., 2007). Of note, acute treatment of VTA slices with estrogen receptor antagonists did not decrease basal firing rates of VTA DA neurons. We also did not observe a difference in baseline firing rates in OVX mice treated with E2 compared with VEH. It is possible that the slightly higher firing rate observed in estrus compared with diestrus is not due to ongoing estrogen receptor activity, but instead might result from actions of progesterone or progesterone-derived neurosteroids (Maguire et al., 2005).

The potential signaling mechanism(s) by which ERα increased the ethanol sensitivity of VTA DA neurons was provided by our experiments using an mGluR1 antagonist, which decreased ethanol stimulation of neurons from mice in diestrus and of neurons from OVX mice treated with E2. These results indicate that mGluR1 is required for E2-mediated enhancement of ethanol-induced excitation of VTA DA neurons and suggests that the effects of E2 may be due to rapid signaling at the cell membrane. Estrogen receptors (both ERα and ERβ) functionally couple to mGluRs in various brain regions such as the hippocampus, striatum, and hypothalamus (Dewing et al., 2007; Grove-Strawser et al., 2010; Huang and Woolley, 2012; Oberlander and Woolley, 2016). ERα and mGluRs are associated in complexes at the cell membrane with caveolin proteins (Razandi et al., 2002; Boulware et al., 2007; Dewing et al., 2007; Boulware et al., 2013; Meitzen et al., 2013; Tabatadze et al., 2015; Pastore et al., 2019). Our results support the possibility that a similar coupling of ERα and mGluR1 occurs in the VTA of females and that this interaction is an important regulator of the sensitivity of DA neurons to ethanol excitation. Dissecting the signaling mechanisms downstream of the ERα/mGluR1 interaction involved in ethanol sensitivity will be an important area for future research. For example, in the hippocampus, E2 acts through mGluR1 to increase the production of the endocannabinoid anandamide, which functions in a retrograde manner to activate presynaptic cannabinoid (CB1) receptors, resulting in a suppression of presynaptic GABA release (Huang and Woolley, 2012). This mechanism could conceivably increase the sensitivity of VTA DA neurons to ethanol. Activation of ERα/mGluR1 by E2 in the hippocampus increases phosphorylation of ERK, which is important for the enhancement of memory by E2 (Boulware et al., 2013).

Estrogen receptors in the VTA of female mice are important for promoting binge-like alcohol drinking, as demonstrated by shRNA-mediated knockdown of ERα and ERβ. We found a stronger decrease in ethanol drinking by reducing levels of ERα compared with ERβ. It is possible that this is because ERα, and not ERβ, enhances excitation of VTA DA neurons by ethanol. We observed greater expression of ERα *vs.* ERβ in the VTA, and it is also possible that the increased abundance of ERα is responsible for its more prominent role in alcohol drinking. Reducing levels of ERβ in the VTA modestly decreased binge-like ethanol drinking by females. Although ERβ does not alter the excitatory effect of ethanol on VTA DA neurons, it may play a role in the VTA in alcohol drinking through its actions on other neuronal types or by affecting the innate physiology of VTA DA neurons independently of ethanol stimulation, for instance by altering the expression of various receptors or ion channels, or by altering the physiology of non-DA neurons that indirectly affect DA VTA neurons. Future experiments will examine how ERβ acts to alter neuronal activity in the VTA.

One might have predicted from our electrophysiology results that: 1) ethanol consumption during diestrus would be higher than in estrus because of the greater response to ethanol during this phase, and 2) knockdown of ERα in the VTA would only decrease ethanol drinking during diestrus, because the ERα antagonist MPP only altered ethanol-induced excitation of VTA DA neurons in diestrus and not in estrus. A close examination of our data indicated that there was no effect of estrous cycle phase on alcohol consumption, consistent with our previous findings and those of others (Roberts et al., 1998; Ford et al., 2002b; Priddy et al., 2017; Satta et al., 2018a). Knockdown of ERα in the VTA also did not decrease drinking only during diestrus; ethanol consumption was reduced by knockdown of ERα regardless of estrous cycle phase. Estrogen receptors in the VTA are important for promoting ethanol drinking in female mice, possibly through multiple mechanisms, including enhancing ethanol excitation of VTA DA neurons.

Interestingly, knockdown of estrogen receptors in the VTA of male mice did not affect binge-like ethanol drinking, despite the fact that males express ERα and ERβ in the VTA. Sex differences in the responses to E2 are not unprecedented. For example, E2 suppresses inhibitory neurotransmission in the hippocampus in females, but not in males, and this effect is mediated by ERα that are coupled to mGluR1 (Huang and Woolley, 2012) and the ability of E2 to increase the interaction of ERα and mGluR1 in females but not in males (Tabatadze et al., 2015). Given our results demonstrating that E2 enhancement of ethanol excitation of VTA DA neurons is mediated by a potential ERα/mGluR1 interaction, the same mechanism may be operative in the sex-specific role of estrogen receptors in the VTA on ethanol drinking. A more complete understanding of the E2-dependent signaling mechanism(s) in the VTA that drive binge ethanol drinking is an important area for future research and may lead to more effective treatments to reduce excessive drinking by women.

## Acknowledgements

This work was supported by the National Institute on Alcohol Abuse and Alcoholism (Grant P50 AA022538 to AWL and MSB and grant U01 AA020912 to AWL) and the National Institute on Drug Abuse (Grant R01 DA033429 to AWL).

## Extended data

**Figure 6-1.**
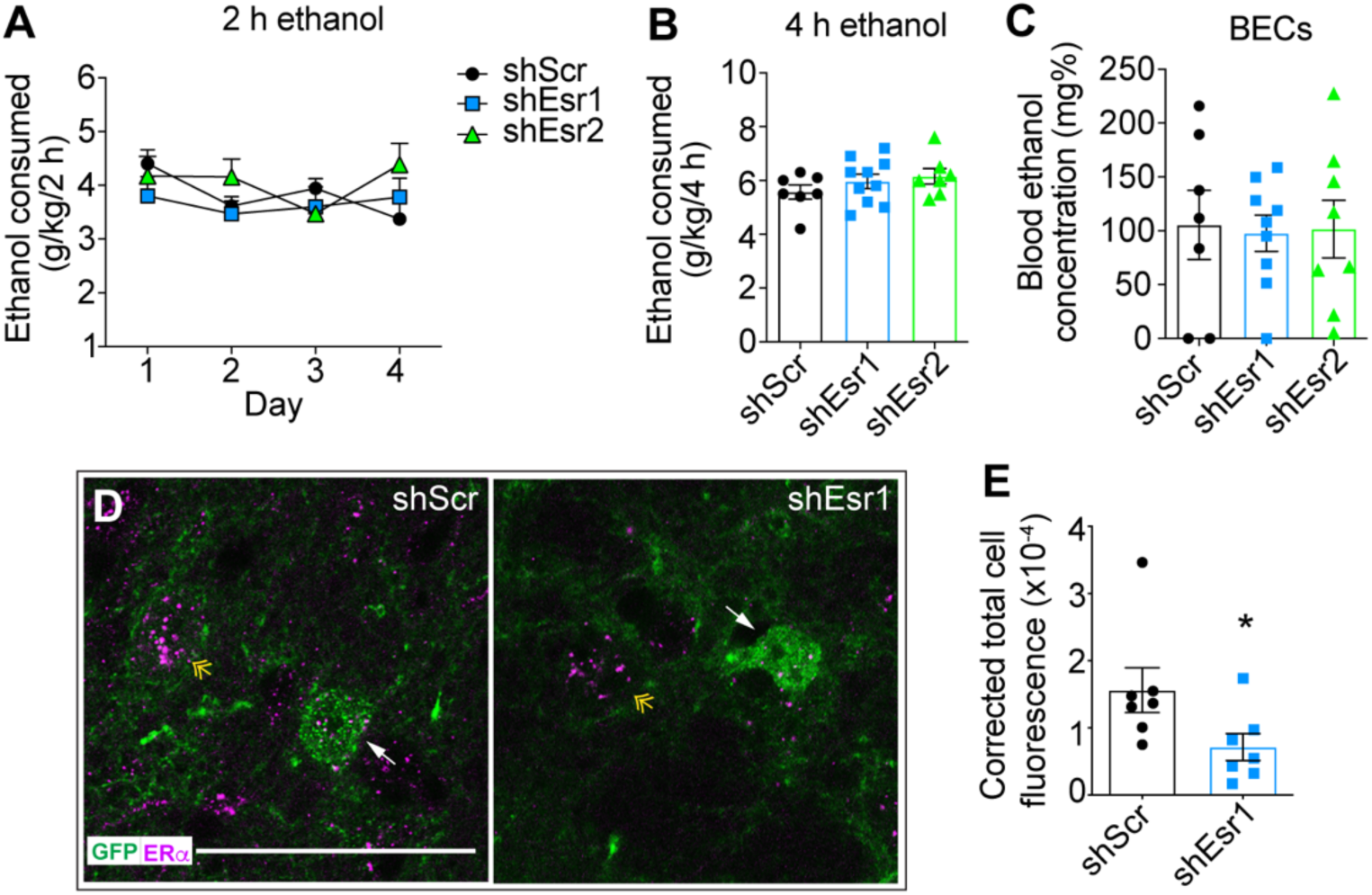
Intra-VTA knockdown of estrogen receptors does not affect binge ethanol consumption in male mice. Gonadally intact male mice were injected with lentivirus expressing either shScr (n=7), shEsr1 (n=10), or shEsr2 (n=7) and tested for binge ethanol drinking in the drinking in the dark test. ***A***, Ethanol intake in g/kg over 2 h on each day of drinking. ***B***, Ethanol intake in g/kg during the final 4-h drinking session. ***C***, Blood ethanol concentrations (BECs) after the final 4-h drinking session. ***D***, Representative images of GFP (green) and ERα (magenta) immunostaining in the VTA after the completion of the drinking test in mice that received lentivirus expressing shScr (right panel) or shEsr1 (left panel). White arrows indicate a GFP-labeled cell with ERα staining in the shScr sample and reduced ERα staining in the shEsr1 sample. Yellow double arrowhead shows an uninfected ERα positive cell. ***E***, Quantification of ERα fluorescence intensity in GFP-positive cells in the VTA of mice receiving virus expressing shScr or shEsr1. Data are presented as corrected total cell fluorescence (calculated as integrated density-area of selected cell x mean fluorescence of background readings). For each group, mean fluorescence from 7 cells (from 5 animals) is shown. \**p*=0.051.

